# A multidimensional analysis of the risk of infection with *Ehrlichia canis* among urban dogs in Iquitos, Peru

**DOI:** 10.1101/2025.07.16.663537

**Authors:** Cusi Ferradas, Oliver A. Bocanegra, Daniela A. Condori, Diego B. Cuicapuza, Fabiola Diaz, Janet Foley, Andrés G. Lescano, Maureen Laroche

## Abstract

*Ehrlichia canis* is a tick-borne bacterium that causes a potentially fatal disease in dogs called Canine Monocytic Ehrlichiosis. In this cross-sectional study, we used a One Health framework to identify statistical associations between *E. canis* infection in dogs and multiple dog-related, human and environmental factors in Iquitos, Peru. Due to the lack of consensus regarding the positivity threshold for *E. canis* qPCR assays, we also evaluated if the factors associated with infection remained conserved regardless of the Ct value cut-off used: Ct<35, a conservative but commonly accepted Ct *cut-off* for bacterial screening, or Ct ≤40, which has been used in several *E. canis* studies. Under the more conservative scenario, we found that the prevalence of *E. canis* among dogs was 19.6% (95% CI 15.8–23.9%). Additionally, we showed that risk factor analyses utilizing a qPCR Ct cut-off of 35 or 40 (with conventional PCR confirmation for samples with a Ct>35) yield comparable results in statistical models, although some differences should be considered. Our findings suggest that in Iquitos, Peru, interventions to prevent *E. canis* infection should prioritize dogs living in houses with corrugated iron walls. Additionally, comprehensive strategies targeting dogs that have recently traveled and incorporating neutering/spaying and widespread acaricide programs may also prove beneficial. We also discuss the challenges encountered during molecular testing for *E. canis* detection, highlighting the broader difficulties of studying poorly understood intracellular pathogens in Global South countries.

## 1. Introduction

Canine monocytic ehrlichiosis (CME) is a multisystemic disease caused by *Ehrlichia canis*, an obligate intracellular pathogen, whose tick vectors belong to the *Rhipicephalus sanguineus* complex. *Ehrlichia canis* is distributed worldwide, but is more prevalent in the tropics and subtropics, sharing distribution with *Rhipicephalus linnaei* (previously known as *R. sanguineus* tropical lineage), its local vector. Human ehrlichiosis, caused by *E. canis,* is rare but associated with potentially severe disease, with symptoms that include fever, headaches, malaise, leukopenia, thrombocytopenia, renal failure, and elevated liver enzymes ^1,2^.

Wild canids and domestic dogs (referred to as “dogs” hereon) are susceptible to *E. canis* infection. In dogs, the course of the disease starts with an incubation period of 8 to 20 days followed by the acute phase, which lasts two to four weeks. During the acute phase of the disease, dogs can present mild to moderate nonspecific signs, such as fever, depression, lethargy, and anorexia. However, systemic infection can result in more severe complications such as hemorrhages, blindness, and neurological manifestations ^3^. Some dogs recover from the acute form of the disease, either spontaneously or with treatment, while others develop the subacute phase of the disease, which can last from months to years ^4^. A subset of dogs in the subacute phase transitions into the chronic phase of the disease which can last for months or years. Dogs with chronic CME can demonstrate bone marrow hypoplasia, which is generally unresponsive to treatment and results in a high mortality rate ^5^. The potential for fatal outcomes of CME among dogs emphasizes the need to conduct studies that expand the understanding of the local epidemiology of the disease and support improved preventive and control strategies.

In Global South countries*, various methods are used for the diagnosis of *E. canis*, including initial clinical diagnosis (based on symptoms and hematological abnormalities), microscopic examination of *E. canis* morulae in Giemsa-stained blood, serological diagnostic tests (including rapid tests, such as the SNAP 4Dx (IDEXX, Westbrook, ME)) and, rarely, molecular diagnostic techniques, such as polymerase chain reaction (PCR). Serological tests may indicate previous infection or cross-react between *Ehrlichia* species^3^. In contrast, molecular techniques can detect active infection and be species-specific. Real-time PCR (qPCR) has higher sensitivity and specificity than conventional PCR, allowing bacterial detection despite the low bacteremia associated with early or late stages of the disease ^6^. The cycling threshold (Ct) cutoff value below which *E. canis* qPCR assays are each considered positive often varies across laboratories. *Ehrlichia canis* is often associated with low bacterial loads at the time of testing, thus, some researchers use higher cut-offs – as high as 40 – to improve detection sensitivity, at the potential cost of specificity ^7,8^. Indeed, high Ct values may occur due to probe degradation and/or non-specific amplification and generate false positive results. False positives may then influence the estimated disease prevalence in population-level studies, as well as the detection and strength of associations between exposure variables and disease status. In the absence of an optimized gold standard for detecting *E. canis*, it is important to compare models with varying cut-offs to determine whether the identified infection risk factors remain consistent despite these molecular biology discrepancies.

In Iquitos, Peru, a high seroprevalence of CME has been reported among dogs with compatible clinical signs, yet, no studies have applied molecular diagnostic techniques which would allow accurate and timely characterization of the acute infection and, therefore, a sensitive evaluation of risk factors ^9^. Iquitos City is the capital of the Department of Loreto, located in the Peruvian Amazon region, in the Northeast of Peru. It is a landlocked city, only accessible by boat from its riverbed and by air. The city and houses are characterized by a precarious infrastructure, and in 2018, 8.6% to 26.5% (median: 20.35%) of the individuals from the four districts forming Iquitos City – Iquitos, Punchana, Belén, and San Juan – lived in poverty ^10^. Although no studies have estimated the dog population in Iquitos City (referred to as “Iquitos” hereon), it is known by local veterinarians that high numbers of stray and community (owned, free-ranging) dogs are roaming across the city (Personal communication, Yauri, 2023). Previous studies on CME in the Americas have found that risk factors for *E. canis* infection vary by geographic area ^11–13^. This variation may be due to the diverse house characteristics, dog behavior, cultural customs related to attitudes towards dogs, as well as other factors related to hosts and vectors. A comprehensive understanding of the risk factors for infection with *E. canis* requires a One Health approach that comprehensively examines factors related to dogs, their owner(s), and their environment. Such factors include, for example, dog signalments, owners’ knowledge about ticks and tick-borne diseases, education level, and use of acaricide medication, as well as house wall and floor materials, which can directly impact tick infestations ^14^.

Understanding the burden of *E. canis* infection and the relevant factors associated with CME is crucial to designing effective surveillance and prevention, and targeted control measures, especially in low-resource settings where regular veterinary interventions are lacking. Therefore, the objectives of this study were: 1) to investigate the prevalence of *E. canis* among dogs from Iquitos, and 2) characterize dog-, owner-, and house-related factors associated with *E. canis* infection among dogs living in Iquitos, Peru, under two models with different qPCR positivity thresholds. The first model prioritized higher specificity and was more conservative, considering only samples with Ct < 35 as positive, as this cut-off is often accepted for bacterial diagnosis among the scientific community ^15^. The second model, designed for higher sensitivity, while retaining specificity, considered samples with Ct < 35 and those with Ct between 35 and 40 that were confirmed positive by conventional PCR.

## 2. Methods

### 3.1. Study design

We conducted a cross-sectional study nested in the study “Eco-epidemiological study of ectoparasite-borne diseases in Peru”, which was approved by the Gerencia Regional de Salud (GERESA) Loreto and by the Universidad Peruana Cayetano Heredia (UPCH) IRB and IACUC (project number: 201148).

Sample collection for this study was conducted in Iquitos (Peru) in April 2023.

Iquitos (3.7437° S, 73.2516° W) has a total population of 479,866 individuals (2017 national census) and comprises four districts: Iquitos, San Juan Bautista, Punchana, and Belen. April is in the rainy season, with daily temperatures ranging from 22 to 32 °C. During the rainy season, Iquitos experiences widespread flooding in various areas: Nuevo Versalles and Masusa in Punchana district; riverbank area of the historic center and Moronococha in Iquitos district; Zona baja in Belen district; and Pampachica, Cabo López, and Nueve de Octubre in San Juan district. Zona baja in Belen is the largest and most severely flooded sector. April was chosen to conduct field activities as the 2023 rainy season was ending, thus allowing us to access most areas of the Belen district without the need for a boat, thereby reducing costs, and minimizing the likelihood of fieldwork disruptions due to extreme heat and heavy rainfall.

In the present study, we included dogs living in houses in Iquitos City, which is formed by the urban areas of the four aforementioned districts. Exact district border information was obtained from the Peruvian National Institute of Statistics and Informatics (Figure 1). Accordingly, the following district borders were used: the start of the Iquitos-Nauta highway (southwest), the Nanay river (northwest), the Amazonas river (northeast), and the Itaya river (southeast), as shown in Figure 1.

**Figure 1.**
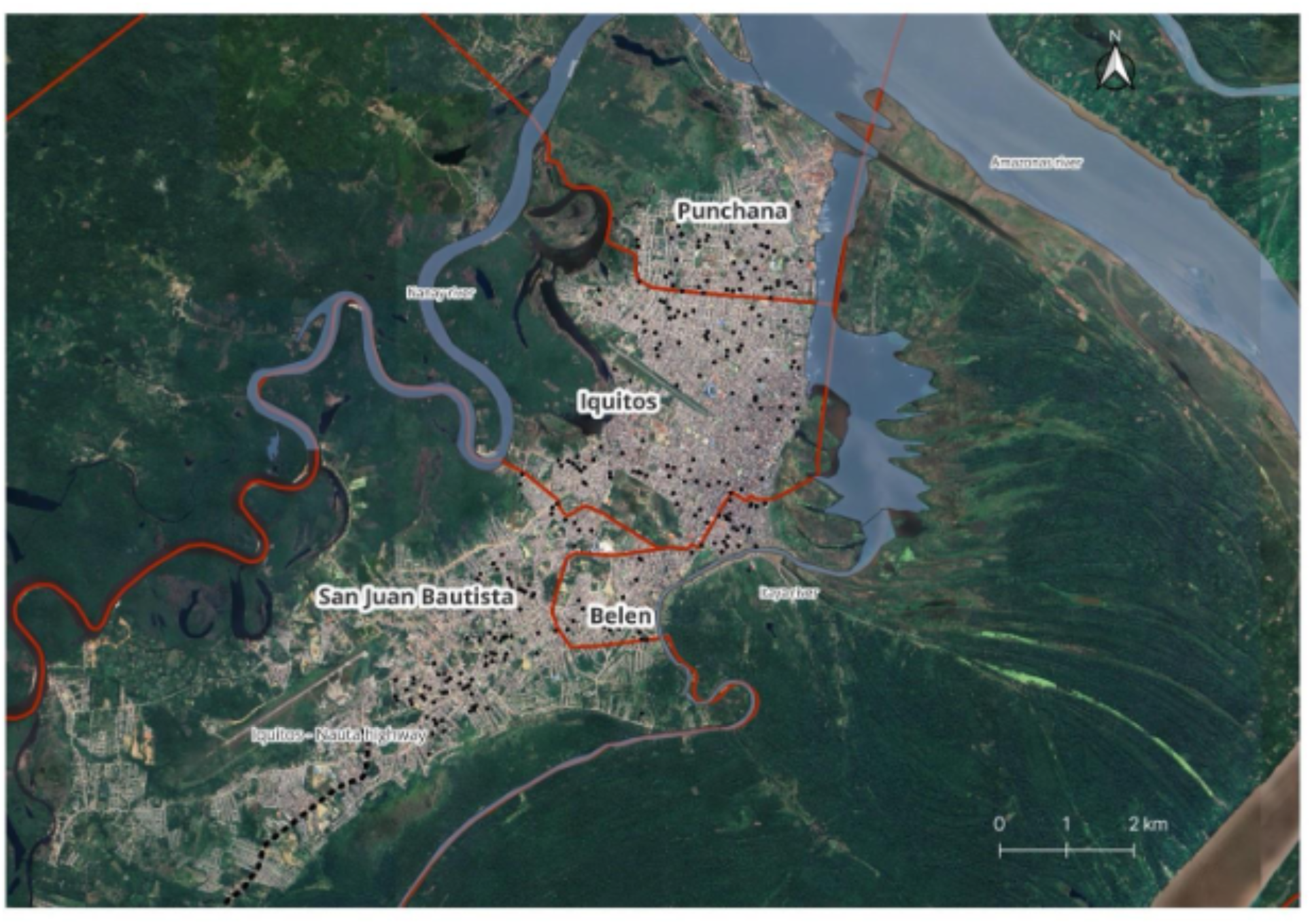
Map of Iquitos City showing the study area’s borders. Borders include the start of the Iquitos-Nauta highway (southwest), the Nanay river (northwest), the Amazonas river (northeast), and the Itaya river (southeast). Homes enrolled in the study are shown as black dots.

### 3.2. Sample size calculation and house selection

A minimum sample size of 286 dogs, equivalent to 286 houses (considering a conservative scenario of one dog per house), was calculated to evaluate factors potentially associated with *E. canis* infection. We chose 80% power and 0.95 confidence level, and assumed a 0.2 intraclass correlation coefficient. We based our sample size calculation on a previous study conducted in Panama, focusing on dogs’ age and veterinary clinic visits as key risk factors ^16^. The calculation was done in Stata 15.0 (Stata Corp., College Station, TX).

We randomly selected 286 buildings in QGIS v.3.28.0-Firenze (https://www.qgis.org) using a base map of all the buildings present in Iquitos city provided by Dr. Amy Morrison (University of California, Davis). According to the 2017 national census, the number of houses in Iquitos (28,381) and San Juan Bautista (30,037) districts is approximately twice the number of houses in Belen (13,467) and Punchana (15,698). Therefore, we randomly selected 96 buildings in Iquitos and San Juan each, and 47 in Belen and Punchana each to ensure adequate representation of each district. The selection was done using the random selection algorithm in QGIS v.3.28.0-Firenze. Then, we used the centroids algorithm to obtain the geographic coordinates (GC) of each house. The centroids GC were downloaded as a *kml* file that was uploaded to the maps.me application (My.com, VK holding). This application was used in the field to locate the selected houses.

In the field, the research team visited each randomly selected building. If the selected building was a business, an abandoned house, or nobody was present, we walked to the right until we found a house with someone present.

### 3.3. House visits, participant recruitment, and questionnaire

Two field teams, each comprising a veterinarian, a veterinary technician, and two biologists, were deployed. Houses with someone older than 18 years old present and at least one dog owned were selected for enrollment, and informed consent was secured by explaining the study procedure, benefits, and risks. Ineligible houses were noted as such, and the research team walked to the next house to the right.

After receiving informed consent from participants, the number of people and dogs in each house were recorded prior to verbally administering a questionnaire to the dog owner or any person from the house involved in the dog’s care (referred all as “owner” hereon). To limit biases, each field team designated a member in charge of consistently administering the questionnaire until the end of the study. Our questionnaire (Sup. material 1) included 5 sections: **owner’s sociodemographic information** (age, gender, education level, occupation, place of work, contact with animals at work, and income level); **house characteristics** (electricity availability; water source; type of sewage system and presence of open sewage nearby; presence of nearby dumps/abandoned houses/food markets, and presence of stray animals around the house**); knowledge, experiences, and risk perception regarding ticks and tick-borne diseases** (if owner was able to identify ticks in different developmental stages, if owner knew ticks feed on various animals and humans, that people and dogs can get tick-borne diseases, including CME, how concerned owners were about tick-borne diseases in dogs and humans, and if they had ever found a tick attached to their bodies); **dog’s epidemiological data** (pet signalment, frequency of veterinary visits, lifestyle, type and frequency of acaricide use, travel and tick infestation during the last year); and **dog clinical data** (presence of clinical signs compatible with CME during the last month, mucous membranes color, body condition, and skin turgor test).

Additionally, we visually inspected the house to determine the main wall and floor material(s) inside the house and in the patio (if the house had one), recording presence or absence of each type of wall or floor material.

The exhaustive list of assessed variables is available as Supplementary Data.

### 3.4. Sample collection

A veterinary technician (accompanied by a veterinarian) collected blood samples from a maximum of two dogs in the house. If more than two dogs lived in the house, we randomly selected two of them (using SocialGest, https://www.socialgest.net/es/sorteos-gratis for sample randomization). We only sampled dogs if their owners indicated that they were not aggressive. Dogs were safely muzzled and manually restrained. A maximum of 6 ml of blood was collected from dogs in vacuum blood collection tubes with EDTA.

All blood samples were immediately stored on ice and a cold chain was maintained until they were processed at the Emerging Diseases and Climate Change Research Unit (Emerge) laboratory in UPCH, Lima, Peru.

Ticks were also conveniently collected from dogs to be identified and to improve the well-being of canine participants. Dogs were systematically inspected from head to tail. Special attention was paid to the ears, neck, between the front and back legs, between the toes, and around the tail. Additionally, house walls and floors were visually inspected to ascertain environmental tick infestation and collect any ticks found. Collected ticks were placed in cryovials containing 70% ethanol and kept at room temperature at UPCH. A maximum of 20 ticks were placed in the same cryovial if they were completely covered by ethanol. One cryovial was used for each dog and ticks collected from their house were stored in another, with a specific labeling code linking all cryovials from the same location.

### 3.5. DNA extraction and real time-PCR

DNA was extracted from canine whole blood samples using the DNeasy Blood and Tissue kit (Qiagen, Valencia, CA) following the kit’s protocol. We added 3 µl of an internal amplification control (IAC) before the addition of the AL buffer. The IAC is a linearized primer containing a sequence of *Arabidopsis thaliana* and was added to identify potential qPCR false negative results.

A previously standardized qPCR targeting the *E. canis* disulfide bond formation protein gene (*dsb*) was performed using primers and probes shown in **Sup. Table 1** and the following thermocycling conditions: 50 °C for 2 min, 95 °C for 10 min, and 50 cycles of 95 °C for 15 seconds, followed by 60 °C for 1 minute^6,12^. A *dsb* gene gblock gene fragment was used as the positive control (IDT, IA, USA). Water was used as the negative control. Samples with a qPCR cycle threshold (Ct) below 35 were considered positive.

**Table 1.** House-related factors: House characteristics in Iquitos city. *Wall and floor materials shown in this table indicate that at least some part of the walls or floors were made of the material indicated. Total house number: n=285.

|  | N (%) |
| --- | --- |
| **District of residency** |  |
| Iquitos | 96 (33.7) |
| San Juan | 95 (33.3) |
| Punchana | 47 (16.5) |
| Belén | 47 (16.5) |
| **Wall material\*** |  |
| Brick or cement | 242 (84.9) |
| Plywood | 147 (51.6) |
| Wood | 130 (45.6) |
| Corrugated iron | 61 (21.4) |
| **Floor material\*** |  |
| Cement | 217 (76.1) |
| Tile | 86 (30.2) |
| Dirt | 59 (20.7) |
| Wood | 24 (8.4) |

### 3.6. Conventional PCR and Sanger sequencing

Samples that had a qPCR Ct value between 35 and 40 were run by conventional PCR targeting a segment of the Tandem Repeat Protein 36 gene (*trp36*) to confirm positivity, using the TRP36-F2 and TRP36-R1 (**Sup. Table 1**) ^17^. All conventional PCR were performed in 25 μl reaction volumes containing 3.5 μl of nuclease-free water, 2.5 μl of each primer (at a 10uM concentration), 12.5 μl of GoTaq Green Master Mix 2X (Promega, Madison, WI), and 4 μl of DNA. Amplification was carried out by denaturation at 95 °C for 5 min, 35 cycles of denaturation (30 s, 95 °C), annealing (1 min, 50 °C), and extension (1 min, 72 °C), and a final extension of 72 °C for 5 min. This protocol was optimized from a previously published protocol^17^. A sample with a qPCR Ct of 24.33 was used as a positive control, while a qPCR negative sample and a blank sample (product of DNA extraction without the addition of a sample) were used as the negative controls.

Conventional PCR amplicons from three samples with a Ct lower than 35 and 12 samples with a Ct between 35 and 40 were sequenced. These samples were amplified using 6 μl of nuclease-free water, 1.25 μl of each primer (at a 10uM concentration), 12.5 μl 2X Phusion Master Mix Buffer (Thermo Scientific, Waltham, MA), and 4 μl of DNA. Amplification was carried out by denaturation at 98 °C for 1 min, 35 cycles of denaturation (10 s, 98 °C), annealing (30 sec, 54.6 °C), and extension (30 sec, 72 °C), and a final extension of 72 °C for 5 min. The 12 samples with a Ct between 35 and 40 that were processed for sequencing, were amplified using the GoTaq Green Master Mix 2X as previously described while the three samples with Ct < 35 were amplified with a Phusion High Fidelity Taq DNA polymerase.

These 15 samples were sent to Macrogen Inc., Chile for purification and Sanger sequencing on the 3730xl DNA Analyzer platform (Applied Biosystems). Sequence evaluation and alignment were performed using Molecular Evolutionary Genetics Analysis (MEGA X). Sequences were compared between them and those in NCBI GenBank using the Basic Local Alignment Search tool (BLAST).

### 3.7. Statistical analysis

The covariates included in the analysis, defined a priori, were organized as dog-, house-, and owner-related factors (**Sup. Table 2)**.

**Table 2.**
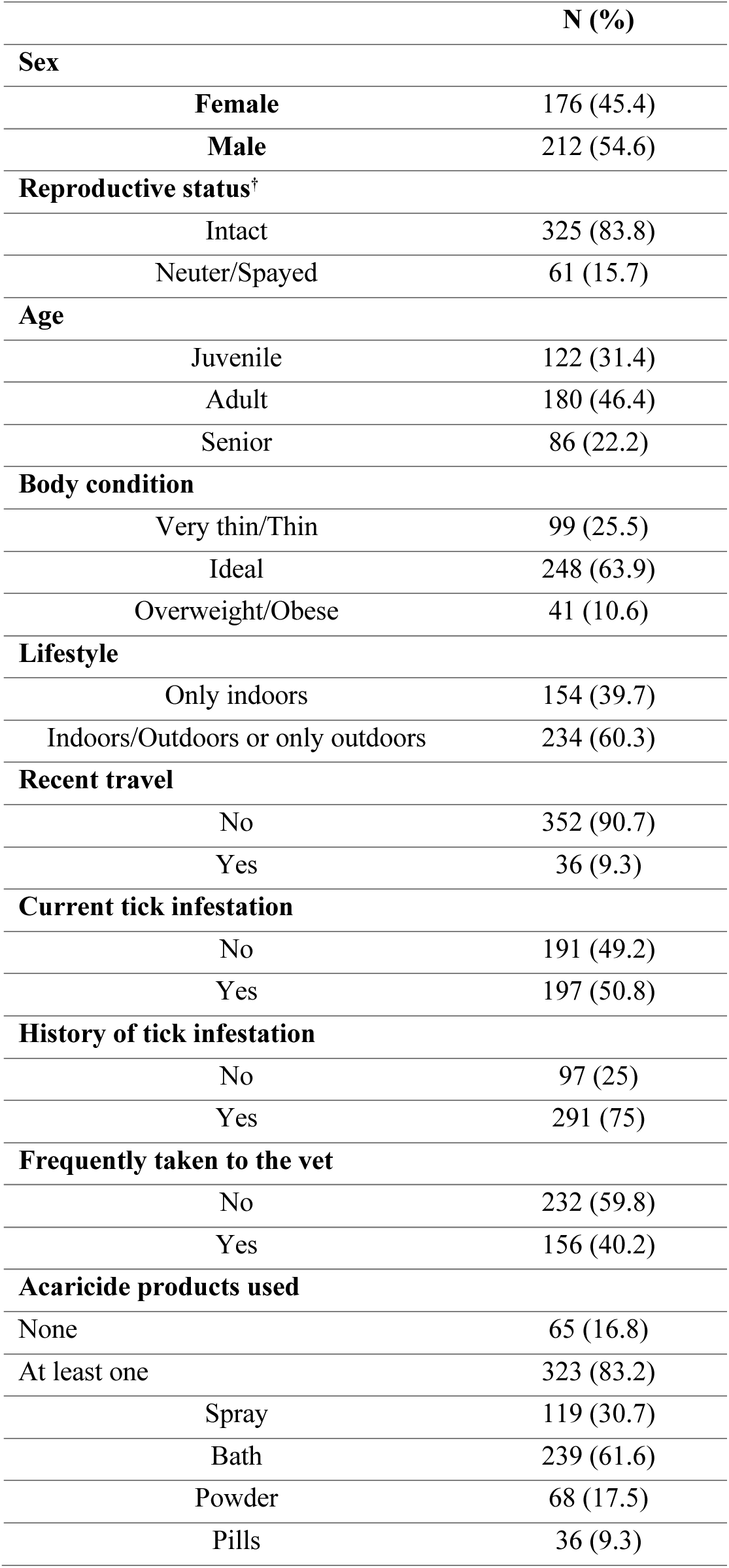
Dog-related factors: Dog characteristics and veterinary care in Iquitos city. ^†^Percentages may not sum up to 100% due to missing values. Total number of dogs: n=388.

|  | N (%) |
| --- | --- |
| **Sex** |  |
| Female | 176 (45.4) |
| Male | 212 (54.6) |
| **Reproductive status<sup>†</sup>** |  |
| Intact | 325 (83.8) |
| Neuter/Spayed | 61 (15.7) |
| **Age** |  |
| Juvenile | 122 (31.4) |
| Adult | 180 (46.4) |
| Senior | 86 (22.2) |
| **Body condition** |  |
| Very thin/Thin | 99 (25.5) |
| Ideal | 248 (63.9) |
| Overweight/Obese | 41 (10.6) |
| **Lifestyle** |  |
| Only indoors | 154 (39.7) |
| Indoors/Outdoors or only outdoors | 234 (60.3) |
| **Recent travel** |  |
| No | 352 (90.7) |
| Yes | 36 (9.3) |
| **Current tick infestation** |  |
| No | 191 (49.2) |
| Yes | 197 (50.8) |
| **History of tick infestation** |  |
| No | 97 (25) |
| Yes | 291 (75) |
| **Frequently taken to the vet** |  |
| No | 232 (59.8) |
| Yes | 156 (40.2) |
| **Acaricide products used** |  |
| None | 65 (16.8) |
| At least one | 323 (83.2) |
| Spray | 119 (30.7) |
| Bath | 239 (61.6) |
| Powder | 68 (17.5) |
| Pills | 36 (9.3) |

We conducted two statistical analyses with different outcomes to assess whether the associated risk factors remained consistent regardless of the Ct cut-off used: i) samples with a qPCR Ct < 35 were considered positive, ii) qPCR positive samples with a Ct ≤ 40 and confirmed by conventional PCR were considered positive for *E. canis*.

We evaluated the association between a set of covariates, shown in the conceptual framework (Figure 2), and both outcomes. We used mixed effects logistic regression models (melogit command on Stata) with a random intercept to account for clustering of dogs at the house level. Bivariate analyses were conducted to evaluate the association between *E. canis* positivity and dog-, house-, and owner-related factors for each aforementioned outcome.

**Figure 2.**
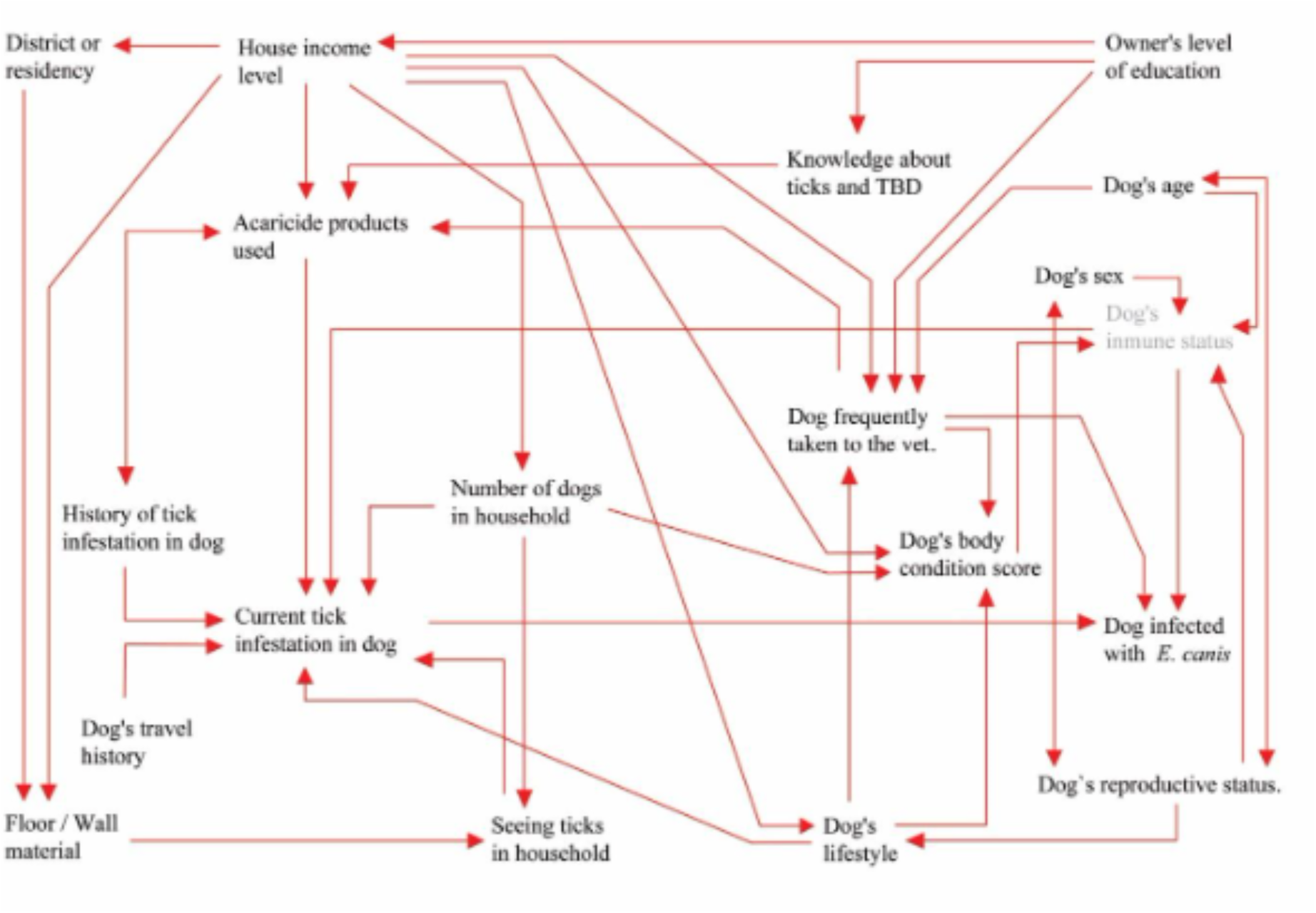
Conceptual framework of factors hypothesized to be associated with *E. canis* infection among dogs living in Iquitos. Arrows indicate potential associations between the factors.

Manual forward nested models were built to determine which variables entered the final multilevel multivariable regression models of factors associated with *E. canis* qPCR positivity. Here, variables were sequentially added and retained only if their inclusion resulted in a decrease in the model’s Akaike Information Criterion (AIC), which balances model fit and complexity. A total of 61 observations were excluded from this model due to missing values in the covariates (the total number of observations included was 327, which belong to 243 houses). Because 39/61 (63.9%) missing values were due to owners’ unwillingness to share their house income, we compared all factors included in the analysis as covariates between individuals who did not indicate their house income and individuals who did indicate their house income (**Sup. Table 3**). For the dog-related factors, one dog from the house was randomly selected. Multicollinearity between variables included in the final model was assessed using the Variance Inflation Factor (VIF).

**Table 3.** Owner-related factors: Owners’ sociodemographic characteristics and knowledge about ticks and tick-borne diseases in Iquitos city. ^†^Numbers may not sum up to the total number of dogs included in the study due to missing values. Total number of owners: n = 285.

|  | N (%) |
| --- | --- |
| **House income level<sup>†</sup>** |  |
| ≤ S/. 500 | 23 (8.1) |
| S/. 501 – 1,000 | 29 (10.2) |
| S/. 1,001 – 1,500 | 47 (16.5) |
| S/. 1,501 – 2,000 | 32 (11.2) |
| S/. 2,001 – 2,500 | 41 (14.4) |
| > S/. 2,500 | 85 (29.8) |
| **Level of education<sup>†</sup>** |  |
| No studies – Elementary school (complete) | 34 (11.9) |
| High school (incomplete) | 24 (8.4) |
| High school (complete) | 78 (27.4) |
| Institute (incomplete) | 26 (9.1) |
| Institute (complete) | 46 (16.1) |
| University (incomplete) | 31 (10.9) |
| University (complete) – Graduate school | 45 (15.8) |
| **Knowledge about ticks and TBDs** |  |
| Common to see ticks in dogs in Iquitos | 226 (79.3) |
| Tick identification | 130 (45.6) |
| Dogs can get TBDs | 242 (84.9) |
| Names TBDs that affect dogs | 23 (8.1) |
| Heard about CME | 48 (16.8) |
| Humans can get TBDs | 195 (68.4) |

Statistical analyses were done using a confidence interval of 95% in Stata 15.0 (Stata Corp., College Station, TX). Variables with a p-value < 0.05 were considered statistically significant.

## 4. Results

### 4.1. House enrollment and characteristics

We visited 714 houses to achieve our target sample size of 286 occupied houses with dogs, where owners agreed to participate in the study. The number of occupants in all visited houses ranged from 1 to 21 (median 4, IQR 3-6), for a total of 3,318 individuals. Dogs were found in 416/714 (58.3%) houses, and the number of dogs per house ranged from 1 to 10 (median 1, IQR-1-2), for a total of 690 dogs. This resulted in an owned dog:human ratio of 0.21.

Participation consent was obtained in 286 (68.8%) of the 416 houses with dogs. In these 286 houses, the number of dogs also ranged from 1 to 10 (median 1, IQR-1-2), with most houses having one (174, 60.8%) or two (72, 25.3%) dogs. A total of 389 dogs were found in these houses; however, we were not able to collect blood samples from one of the dogs, resulting in a total of 388 dogs living in 285 houses enrolled in the study.

Most houses had at least one wall made of brick or cement (242, 84.9%) and some floors made of cement (217, 76.1%). The second most common wall material was plywood (147 houses, 51.6%), while the floor material was tile (86, 30.2%). Other wall materials included wood and corrugated iron, while dirt and wood were also found as floor materials. Detailed characteristics of the 285 houses included in this study are shown in **Table 1**.

### 4.2. Dog characteristics, veterinary care, and clinical signs

We collected samples from juvenile (122, 31.4%), adult (180, 46.4%), and senior (86, 22.2%) dogs. Most of the enrolled dogs were male (212, 54.6%) and not neutered/spayed (325, 83.8%) and had an ideal body condition (248, 63.9%), grading as 3 per the 1-5 body condition score scale of the American Animal Hospital Association. However, 99 (25.5%) of them were very thin or thin. Two hundred and thirty-four (60.3%) of dogs had an indoor/outdoor or only outdoor lifestyle, but only 36 (9.3%) of them had traveled to another area of Iquitos or to/from another department of Peru during the last year (**Table 2**).

Only 156 (40.2%) dogs were regularly taken to a veterinarian (at least once per year), but 323 (83.2%) of them received frequent acaricide treatment to prevent/eliminate ticks. The most frequently used acaricidal approach were baths (239 dogs, 61.6%). Among dogs that were frequently bathed, 73 (61%) and 54 (79%) also received an acaricide in spray or powder, respectively. The most used acaricides were Shampoo Amigo (Amitraz 0.25%), Rey Peluchín spray (Fipronil 0.025%), and the Peluchín powder (Cypermethrin 1 g, Salicylic Acid 1 g, soap base C.S.P 100 g). Only 36 (9.3%) of the dogs received oral tick prevention drugs, and the most used was Bravecto® (Fluralaner 250 mg).

We asked owners if their dogs had had symptoms associated with CME during the last month. A hundred and thirteen (29.1%) had vomited; 51 (13.1%) had had diarrhea; 30 (7.7%) had presented some type of bleeding (two presented mouth bleeding, 19 melena, three hematuria, and 8 presented bleeding in other areas); 73 (18.8%) presented weakness; and 68 (17.5%) presented decreased appetite. *Ehrlichia canis* qPCR positivity was not found to be significantly associated with any of these specific symptoms in our sample (results not shown).

After physical examination, 114 (29.4%) and 91 (23.5%) of the dogs presented pale mucous membranes and increased skinfold time (more than two seconds), respectively. Dogs with pale mucous membranes had significantly higher odds of being *E. canis* qPCR positive (Ct<35) than those with pink mucous membranes in the bivariate analysis (OR 2.74 [95% CI 1.38-5.50], p-value = 0.004). The association between an increased skinfold return and *E. canis* qPCR positivity was not statistically significant (p-value = 0.399).

### 4.3. Owners’ sociodemographic information and knowledge about ticks and tick-borne diseases

The house income levels, and the highest educational level achieved by dog owners were highly variable (**Table 3**).

Regarding knowledge about ticks, 226 (79.3%) of the owners knew that tick infestation in dogs is common in Iquitos, however, only 130 (45.6%) of the owners were able to identify ticks in different developmental stages. Regarding knowledge about tick-borne diseases (TBD), most owners (242, 84.9%) knew that ticks can transmit pathogens to dogs; however, only 23 (8.1%) were able to name at least one TBD or tick-borne pathogen, and only 48 (16.8%) had heard of CME. Finally, 195 (68.4%) owners knew that ticks can also transmit pathogens to humans (**Table 3**).

### 4.4. *Ehrlichia canis* PCR and sequencing

Of the 388 dogs tested for *E. canis* by qPCR, 76 tested positive (qPCR Ct < 35), resulting in a positivity rate of 19.6% (95% CI 15.8 – 23.9%). The median Ct was 32.4 (IQR 30.8-34.03).

Additionally, 69 samples had a qPCR Ct between 35 and 40. Of them, 55 (80%) were confirmed positive by conventional PCR targeting the *trp36* gene of *E. canis*. Therefore, a total of 131 samples were confirmed to be positive for *E canis*, resulting in a final positivity rate of 33.8% (95% CI 29.1 – 38.7%).

Five of 15 the (33.3%) samples sent to Macrogen Inc., Chile were successfully sequenced. Out of these five samples, three were amplified using GoTaq Green Master Mix 2X (Promega, Madison, WI) and the other two were amplified using Phusion Master Mix Buffer (Thermo Scientific, Waltham, MA).

All sequences were identified as *E. canis* with percent identities ranging from 83.02% to 100% and query coverage values ranging from 39.0% to 100%. However, due to the quality of the sequences and limited availability of reagents, robust phylogenetic analysis was not possible during this study.

### 4.5. Factors associated with *E. canis* positivity

#### 4.5.1. First analysis (based on samples with Ct < 35)

In the **bivariate analyses**, of <u>dog-related factors</u>, male dogs (OR 2.34, [95% CI 1.20-4.57], p-value = 0.013) and those who had recently traveled (OR 3.68 [95% CI 1.36-9.98, p-value = 0.010]) had statistically significantly higher odds of being *E. canis* positive (**Sup. Table 4**). In contrast, neuter/spayed dogs had lower odds (OR 0.27, [95% CI 0.08-0.89], p-value = 0.031) of being positive. Of the <u>house-related factors</u>, dogs living in houses with at least some walls made of corrugated iron (OR 2.76, [95% CI 1.26-6.05], p-value = 0.011) had higher odds of testing positive, and none of the <u>owner-related factors</u> were found significantly associated with *E. canis* infection.

**Table 4.** Multivariable analysis of factors associated with *E. canis* positivity (qPCR Ct<35) *Mixed effects logistic regression with a random intercept to account for clustering of dogs at the house level. Reference categories are listed in Table 1 and Sup. Table 1. **Wall and floor materials shown in this table indicate that at least some part of walls or floors were made of the material indicated.

|  | OR* | 95% CI | p-value |
| --- | --- | --- | --- |
| **Dog-related factors** |  |  |  |
| Male | 1.89 | 0.90-3.96 | 0.091 |
| Recent travel | **3.59** | **1.19-10.89** | **0.024** |
| Neuter/spayed reproductive status | 0.27 | 0.07-1.05 | 0.058 |
| **House-related factors** |  |  |  |
| Wall material – Corrugated iron** | **2.92** | **1.22-6.96** | **0.016** |
| **Owner-related factors** |  |  |  |
| Common to see ticks in dogs in Iquitos | 2.88 | 0.97-8.55 | 0.056 |

Variables from all three groups (dog-, house, and owner-related factors) were retained in the adjusted model after performing the manual forward nested model analysis (**Table 4**). In the **multilevel multivariable regression models** (**Table 4**), dogs who traveled recently (OR 3.59, [95% CI 1.19-10.89], p-value = 0.024) and dogs living in houses with at least some part of walls made of corrugated iron (OR 2.92, [95% CI 1.22-6.96], p-value = 0.016) had statistically significantly higher odds of being positive to *E. canis* after controlling for the other variables included in the model. While dog’ sex, dog’s reproductive status, and owners’ knowledge that tick infestation among dogs is common in Iquitos were retained in the model, they were not significantly associated with the defined outcome.

#### 4.5.2. Second analysis (based on samples with Ct ≤ 40 and confirmed by conventional PCR)

In the **bivariate analyses**, of <u>dog-related factors</u>, only dogs’ age and recent travel were significantly associated with being qPCR-positive (**Sup. Table 5**). Adult (OR 2.70, [95% CI 1.03-7.09], p-value = 0.044) dogs and those who traveled recently (OR 3.92, [95% CI 1.11-13.85], p-value = 0.034) also had significantly higher odds of being positive to *E. canis*. Dogs living in houses with at least some walls made of corrugated iron (OR 4.76, [95% CI 1,56-14,54], p-value = 0.006) or brick/cement (OR 3.84, [95% CI 1.00-14.71], p-value = 0.050) and those whose owners knew dogs are commonly infested with ticks in Iquitos (OR 3.23, [95% CI 1.07-9.74], p-value = 0.038) had higher odds of testing positive for *E. canis*.

**Table 5.** Multivariable analysis of factors associated with *E. canis* positivity (positive qPCR and conventional PCR)^⊥^. ^⊥^qPCR Ct<35 plus samples with 35<Ct≤40 positive by conventional PCR *Mixed effects logistic regression with a random intercept to account for clustering of dogs at the house level. Reference categories are listed in Sup. Table 1. **Wall and floor materials shown in this table indicate that at least some part of walls or floors were made of the material indicated.

|  | OR* | 95% CI | p-value |
| --- | --- | --- | --- |
| **Dog-related factors** |  |  |  |
| Male | 1.79 | 0.81-3.95 | 0.149 |
| Recent travel | 3.45 | 0.87-13.64 | 0.077 |
| **House-related factors** |  |  |  |
| Wall material – Corrugated iron** | **7.01** | **1.98-24.80** | **0.003** |
| Wall material – Plywood** | 0.52 | 0.21-1.30 | 0.164 |
| **Owner-related factors** |  |  |  |
| Common to see ticks in dogs in Iquitos | 3.06 | 0.95-9.83 | 0.061 |

In the **multilevel multivariable regression models** (**Table 5**), dogs living in houses with at least some part of walls made of corrugated iron (OR 7.01, [95% CI 1.98-24.80], p-value = 0.003) had statistically significantly higher odds of being positive to *E. canis*. Some other variables identified in the bivariate analysis dog-(sex and recent travel), house-(at least some part of the walls made of plywood) and owner-(know dogs are commonly infested with ticks in Iquitos) related factors were retained in the manual forward nested model, indicating that they accounted for some of the outcome variability, but none of them were statistically significantly associated with *E. canis* positivity among dogs.

### 4.7. Tick infestation

Of the 285 owners who answered the questionnaire, 147 (51.6%) indicated seeing ticks in their houses during the last year and a half. We collected ticks from the floor or walls (indoors or outdoors) of 39 houses (13.7%).

Of the 388 dogs sampled, 197 (50.8%) were infested with ticks and, according to owners’ report, 291 (75%) were infested with ticks during the previous year. All ticks were morphologically identified as *R. sanguineus* s.l. Bivariate and multivariate analyses did not show any statistically significant association between current and history of tick infestation and *E. canis* positivity.

## 5. Discussion

Using a One Health framework and molecular techniques, we showed that several variables related to the dogs, their houses, and their owners affect their likelihood of *E. canis* infection in Iquitos, Peru.

The PCR prevalence of *E. canis* among apparently healthy owned dogs was 19.6% based on samples with qPCR Ct < 35, and 33.8% when confirmed samples with Ct from 35-40 were included. Because this is the first report of *E. canis* prevalence in Iquitos, we cannot compare our estimate with previous years. Various studies from Latin America have reported *E. canis*, prevalence rates ranging from 15.3% to 53.6%, however, significant methodological variability exist across studies (e.g., study populations, use of conventional, nested PCR or qPCR, chosen Ct cut-off values, etc. ^12,18–22^). In addition, some discrepancies may be attributed to differences in *R. sanguineus* s.l. infestation prevalence, environmental characteristics, dog ecology and behavior, and the level of care provided by owners for their dogs. Nevertheless, the positivity rate estimated in this study with a qPCR Ct < 35 is comparable to the prevalence estimated in Northern Colombia (15.3%) using the same qPCR assay, but with a Ct cut-off of 40 ^12^.

In our study, recent travel was significantly associated with *E. canis* infection in dogs within the conservative model. Most of these dogs had traveled to rural areas, with others visiting small urban or peri-urban locations (as reported by the interviewees). Since *E. canis* prevalence in the travelled areas is unknown, we are unable to hypothesize why dogs had higher odds of testing positive for *E. canis* after visiting them. **Neutered or spayed dogs** seemed marginally less likely to test positive for *E. canis,* which could stem from the reduced likelihood of sterilized dogs associating in larger dog groups (where infected *R. sanguineus* s.l. can go from one dog to another), a behavior often triggered by the presence of a female in heat. Notably, only a modest 15.7% of dogs were sterilized ^16^. This may be due to owners’ lacking the resources to take their dogs to the veterinarian, their lack of knowledge about the possibility of neutering/spaying, and/or personal beliefs against neutering/spaying (especially for male dogs). Contrary to a previous study, **dog age** was not statistically significantly associated with *E.canis* infection ^16^. Although adult dogs had higher odds of being positive to *E. canis*, the median Ct value among juvenile dogs was the lowest. These results are consistent with repeated exposure and some herd immunity among older dogs, as well as presence of older dogs with subacute or chronic disease, usually associated with lower bacterial loads ^4^.

Neither current tick infestation nor a history of tick infestation (reported by the owner) was associated with qPCR positivity for *E. canis* in either model. The association between tick infestation and PCR positivity has varied across published studies, with some reporting a significant association and others finding no such link ^19,23^ These disparities are unsurprising as CME is a disease that can progress into chronicity, with dogs remaining positive by PCR for extended periods of time, making it extremely challenging to assert that infected ticks collected from an infected dogs were definitively the source of infection. It might therefore be misleading to tentatively correlate tick infestation and qPCR positivity in dogs.

In our study population, although most dogs had an indoor/outdoor lifestyle during the day, 94.59% (367/388) spent nights inside the house. Previous studies have identified that house building materials influence the presence and abundance of various arthropod vectors (triatomines, mosquitos, and sand flies) indoors ^24–26^. In Urabá, Colombia, dirt floors were significantly associated with house infestation with *R. sanguineus* s.l ^27^. In the present study, the odds of *E. canis* positivity were statistically significantly higher in houses with at least part of walls made of corrugated iron. This may be related to corrugated iron generating suitable temperature and humidity conditions inside the house for the development and survival of *R. sanguineus* s.l ^28^. It is important to note that among the 90 houses with corrugated iron incorporated into their walls, the proportion of walls constructed with this material was relatively small: one (0.26%), 67 (17.4%), and 22 (5.71%) houses had 5%, 10%, and 20% of their walls composed of corrugated iron, respectively. We explored if the association between at least part of walls made of corrugated iron and *E. canis* positivity among dogs was due to correlation between presence of this material and another measured factor (other construction materials and house income), but these variables were not correlated in our data. Therefore, it is possible that the presence of walls made of corrugated iron may be a proxy for other, unmeasured factors. Given the diverse range of wall and floor construction materials used in some houses in Iquitos, operationalizing these variables poses a challenge. Therefore, we opted to include the presence (yes/no) of each wall and floor material rather than creating a variable indicating the primary material. Although not statistically significant, walls partially made of plywood accounted for some of the adjusted model’s variability. Our results suggest that the materials of house’s walls may be important drivers of *E. canis* infection among dogs, likely because of their impact on tick populations. Further studies that restrict inclusion criteria to houses with only one construction material should be conducted to further explore the association between this variable and the odds of *E. canis* infection and understand how different construction materials affect the climatic conditions inside the house and the developmental cycles of *R. sanguineus* s.l. in Iquitos.

It should be noted that the person mainly in charge of caring for the dog was not always present during our interviews, which may have impacted our ability to identify owner-related risk factors. Nevertheless, we identified a marginal association between owner awareness of common tick infestations among the general dog population of Iquitos and *E. canis* infection in their dog. This may suggest free-ranging and stray dogs could serve as a source of infection for owned dogs, and/or that past infection with *E. canis* in owned dogs could contribute to increased awareness regarding ticks in owners. This underscores the need for widespread acaricide treatment among owned and free-ranging dog populations.

It is common for *E. canis* bacterial loads to be low during the subacute phase of the disease, leading to false negative results ^8^. Therefore, some investigators use 37 or even 40 as a cycle cut-off for the detection of *E. canis* ^32–34^, a cut-off much higher than what is often accepted for bacterial diagnosis among the scientific community ^15^. Our two scenarios for considering samples positive (qPCR Ct < 35 or Ct between 35 and 40 confirmed by conventional PCR) were used to present conservative and robust results, while simultaneously including samples that could provide relevant information for the characterization of risk factors. The selection of a qPCR Ct cut-off to classify a sample as positive can introduce misclassification bias in epidemiological studies. This misclassification is likely to be non-differential, as it affects all individuals similarly, regardless of exposure or outcome status. In our models, most factors and their associations with the outcome remained consistent across different cut-off values. However, reproductive status improved the AIC in the more conservative model, while the presence of plywood walls in the house was retained in the less conservative model. These findings underscore that, despite qPCR being the most reliable diagnostic test for *E. canis* infection, it has limitations that can affect risk factor analysis and potentially influence recommendations for intervention strategies.

We encountered various challenges while standardizing the conventional PCR assay. Although the TRP36-F2 and TRP36-R1primers have been used in previous studies, discrepant annealing temperatures have been reported as well as a lack of consistency regarding which species of *Ehrlichia* can be captured with these primers, since they have been used to target both *E. canis* and *E. minasensis* ^17,35,36^. Since tandem repeat proteins can have different number of repetitions, we observed bands of different molecular weights in our conventional PCR, which were all sequenced and identified as *E. canis*. Finally, as we did not have a positive control from a confirmed culture, samples that were positive by qPCR with a low Ct value had to be used as a substitute. Due to resources and time constraints, including obtaining laboratory reagents in Peru, we were not able to perform quality control and process samples with a Ct < 35 by conventional PCR nor use other qPCR targets for confirmation. These difficulties underscore that, despite being regarded as inexpensive and easy-to-use in Global North countries, basic molecular methods such as conventional PCR can still be challenging to implement in low– and middle-income countries like Peru.

Research studies that produce optimized DNA extraction methods from a variety of samples and robust standard curves for qPCR cycle cut-offs would improve diagnostic standards for *E. canis*. Nonetheless, we recognize these studies are difficult due to the challenges associated with *E. canis* cell culture. Additionally, although conventional PCR has higher validity compared to other diagnostic techniques (such as blood smears and serological tests), it can remain challenging to standardize and yield inconclusive results requiring sequencing confirmation. This issue is particularly important when working with targets of varying molecular weights, such as tandem repeat proteins.

Some limitations of our study warrant consideration. The cross-sectional study design precludes the assessment of temporality between exposure factors and *E. canis* infection. While we identified active *E. canis* infection using molecular techniques, due to the prolonged course of this disease, the infection may have occurred prior to exposure to certain identified factors. Additionally, certain covariates, such as dogs’ recent travel, lifestyle, and tick infestation during the last year, were self-reported by owners, introducing potential wish and recall bias. Furthermore, our measurement of socioeconomic status and purchasing power for acaricides and veterinary visits was limited to total house income, as the owner’s income was not consistently available during research team visits. Regarding construction materials, we only assessed the presence or absence of wall and floor materials, omitting the quantification of each material percentage due to concerns about subjectivity and potential bias. Future studies should establish standardized methods for quantifying construction materials, as they significantly influence arthropod infestation. The entirety of the fieldwork supporting this study was conducted in April. Given that environmental factors such as humidity and flooding can significantly influence *R. sanguineus* s.l. activity and abundance, future studies should assess whether the prevalence of *E. canis* varies across different seasons and whether these variations can be correlated to *R. sanguineus* s.l. prevalence. Finally, although we collected *R. sanguineus* s.l. ticks, mostly from dogs, we did not test ticks for *E. canis* in this study, as it would have been impossible to distinguish between true tick infection and the presence *E. canis*-infected dog blood within the tick.

In conclusion, this study provides the first assessment of *E. canis* prevalence in dogs and associated factors in Iquitos, Peru. Our results suggest that interventions such as establishing regular, robust campaigns focused on neutering/spaying and widespread acaricide treatment (targeting owned free-ranging and stray dogs) could reduce the incidence of *E. canis* infection and improve the overall health of dogs in Iquitos. Moreover, interventions to prevent tick infestations should prioritize dogs living in houses with at least some walls made of corrugated iron. Our findings and discussion also highlight the challenges of the use of molecular diagnostics in Global South countries. Finally, although some differences should be considered, we showed that risk factor analyses utilizing a conventional qPCR Ct cut-off of 35 yielded comparable results to a statistical model that used a cut-off value of 40 (with conventional PCR confirmation). Further laboratory-based and epidemiological studies should validate the qPCR Ct cut-off for different *E. canis* genes. One of the main limits of the current study design (i.e., cross-sectional) is that it does not allow to determine causality between our different variables. Additional epidemiological studies could provide further insights into factors causally associated with *R. sanguineus* s.l. infestation and *E. canis* infection in this region.

## 7. Acknowledgements

All authors would like to thank our participants, as well as the field team (Brandon Moreno, Magno Zambrano, Carolina Ojeda, Ana del Aguila, Estefany López, Sandra Chávez, and Arturo Chávez) for their dedicated work. Additionally, we would like to thank Dr. Amy Morrison for providing the Iquitos basemap and Román Llontop for his assistance in translating the study questionnaire into English for publication.

## 8. Author contributions

- **Cusi Ferradas:** Conceptualization, data curation, formal analysis, investigation, methodology, project administration, writing-original draft.
- **Oliver A. Bocanegra:** Conceptualization, data curation, formal analysis, investigation, methodology, project administration, writing-review&editing.
- **Daniela A. Condori:** Methodology, investigation, writing-review&editing.
- **Diego B. Cuicapuza-Arteaga:** Methodology, investigation, writing-review&editing.
- **Fabiola Diaz:** Investigation, writing-review&editing.
- **Janet Foley:** Conceptualization, supervision, writing-review&editing.
- **Andrés G. Lescano:** Conceptualization, funding acquisition, methodology, supervision, writing-review&editing.
- **Maureen Laroche:** Conceptualization, methodology, investigation, supervision, writing-review&editing.

## 9. Data availability statement

All data used for this manuscript is available in the main manuscript and its supplementary data.

## 10. Additional informacion

### 10.1. Ethics approval and consent to participate

This study was approved by the Gerencia Regional de Salud (GERESA) Loreto and by the Universidad Peruana Cayetano Heredia (UPCH) IRB and IACUC (project number: 201148). All individuals who participated in the study signed an informed consent.

### 10.2. Consent for publication

All participants consented to the results of this study being published if they did not include information that allowed them to be identified.

### 10.4. Funding

This project was supported by the Fogarty International Center of the National Institutes of Health (NIH) under the Training Grant D43 TW007393 “Emerge: Emerging Diseases Epidemiology Research Training.” The content is solely the responsibility of the authors and does not necessarily represent the official views of the NIH. This project was also supported by the University of California Davis Continuing Graduate Student Internal Fellowship.

### 10.5. Definitions

In this manuscript, inequalities in research are reflected in the use of <u>Global South/Global North</u> countries. Although these terms are imperfect and erase individual national strategies, they reflect with no negative connotation disparities in access to healthcare, resources, and more. In the context of academia, these differences include access to research grants, international conferences, international internships and collaborations, research facilities, high-impact journals, and more.

