## Supplementary data files for "A multidimensional analysis of the risk of infection with *Ehrlichia canis* among urban dogs in Iquitos, Peru"

*Ehrlichia canis* is a tick-borne bacterium that causes a potentially fatal disease in dogs called Canine Monocytic Ehrlichiosis. In this cross-sectional study, we used a One Health framework to identify statistical associations between *E. canis* infection in dogs and multiple dog-related, human and environmental factors in Iquitos, Peru. Due to the lack of consensus regarding the positivity threshold for *E. canis* qPCR assays, we also evaluated if the factors associated with infection remained conserved regardless of the Ct value cut-off used: Ct<35, a conservative but commonly accepted Ct *cut-off* for bacterial screening, or Ct ≤40, which has been used in several *E. canis* studies. Under the more conservative scenario, we found that the prevalence of *E. canis* among dogs was 19.6% (95% CI 15.8–23.9%). Additionally, we showed that risk factor analyses utilizing a qPCR Ct cut-off of 35 or 40 (with conventional PCR confirmation for samples with a Ct>35) yield comparable results in statistical models, although some differences should be considered. Our findings suggest that in Iquitos, Peru, interventions to prevent *E. canis* infection should prioritize dogs living in houses with corrugated iron walls. Additionally, comprehensive strategies targeting dogs that have recently traveled and incorporating neutering/spaying and widespread acaricide programs may also prove beneficial. We also discuss the challenges encountered during molecular testing for *E. canis* detection, highlighting the broader difficulties of studying poorly understood intracellular pathogens in Global South countries.

**Sup. Table 1.** Variables of interest, how they were ascertained and their categories.

| Variable | Ascertained by | Question | Variable measurement | Variable operationalization |
|---|---|---|---|---|
| **Dog-related factors** | | | | |
| Sex | Questionnaire | Is this dog a male or female? | Female<br>Male | Female*<br>Male |
| Reproductive status | Questionnaire | Has this dog been spayed/neutered? | Intact<br>Neutered/Spayed | Intact*<br>Neutered/Spayed |
| Age$^{\gamma}$ | Questionnaire | How old is this dog? | Age in years and months | Juvenile (< 2 years)*<br>Adult (2-7 years)<br>Senior (≥ 7 years) |
| Body condition score$^{\gamma}$ | Physical exam | - | Very thin<br>Thin<br>Ideal<br>Overweight<br>Obese | Very thin/Thin<br>Ideal*<br>Overweight/Obese |
| Lifestyle | Questionnaire | Where do you have this dog most of the time? | Only indoors<br>Indoors and outdoors<br>Only outdoors | Only indoors*<br>Indoors/outdoors or only outdoors |
| Recent travel | Questionnaire | Has this dog traveled recently (other district, province or department)? | No<br>Yes | No, dog did not travel in the last year*<br>Yes, dog traveled in the last year |
| Current tick infestation | Physical exam | - | No, ticks not found<br>Yes, ticks found | No, ticks not found*<br>Yes, ticks found |
| History of tick infestation | Questionnaire | In the last year, have you seen ticks on this dog? | No<br>Yes | No*<br>Yes |
| Frequently taken to the vet | Questionnaire | Do you usually take this dog to the vet? | No<br>Yes | No, dog was not taken to vet in the last year*<br>Yes, dog was taken to vet in the last year |
| Acaricide product used | Questionnaire | Do you use sprays, baths, powder, pills, or other products to eliminate ticks from this dog? | No (to each acaricide)<br>Yes (to at least one acaricide) | None*<br>One or more |
| **Home-related factors** | | | | |
| District | GPS location | - | Iquitos<br>San Juan<br>Punchana<br>Belén | Iquitos*<br>San Juan<br>Punchana<br>Belén |
| Wall material – Brick or cement | Visual inspection** | At least some of the home walls are made of brick or cement | No<br>Yes | No*<br>Yes |
| Wall material - Plywood | Visual inspection** | At least some of the home walls are made of plywood | No<br>Yes | No*<br>Yes |
| Wall material - Wood | Visual inspection** | At least some of the home walls are made of wood | No<br>Yes | No*<br>Yes |

| | | | | |
|---|---|---|---|---|
| Wall material – Corrugated iron | Visual inspection** | At least some of the home walls are made of corrugated iron | No<br>Yes | No*<br>Yes |
| Floor material – Cement | Visual inspection** | At least some of the home floors are made of cement | No<br>Yes | No*<br>Yes |
| Floor material – Tile | Visual inspection** | At least some of the home floors are made of tile | No<br>Yes | No*<br>Yes |
| Floor material – Dirt | Visual inspection** | At least some of the home floors are made of dirt | No<br>Yes | No*<br>Yes |
| Floor material – Wood | Visual inspection** | At least some of the home floors are made of wood | No<br>Yes | No*<br>Yes |
| Number of dogs in the home | Questionnaire | How many dogs lived in this home? | Number of dogs | One*<br>Two<br>Three of more |
| Seeing ticks in the home | Questionnaire | Have you ever seen ticks in your home? | No<br>Yes | No, I have not seen ticks in the last 1.5 years*<br>Yes, I have seen ticks in the last 1.5 years |
| **Owner-related factors** | | | | |
| Home income level | Questionnaire | What is the total monthly income in your home? | S/. 0<br>S/. ≤ 500<br>S/. 501-1,000<br>S/.1,001-1,500<br>S/. 1,501-2,000<br>S/. 2,001-2,500<br>> S/. 2,500 | S/. ≤ 500*<br>S/. 501-1,000<br>S/.1,001-1,500<br>S/. 1,501-2,000<br>S/. 2,001-2,500<br>> S/. 2,500 |
| Level of education | Questionnaire | What is the highest education degree you have completed? | No studies<br>Kindergarten<br>Elementary school (incomplete)<br>Elementary school (complete)<br>High school (incomplete)<br>High school (complete)<br>Institute (incomplete)<br>Institute (complete)<br>University (incomplete)<br>University (complete)<br>Masters/doctorate | No studies – Elementary school (complete)*<br>High school (incomplete)<br>High school (complete)<br>Institute (incomplete)<br>Institute (complete)<br>University (incomplete)<br>University (complete) - Masters/doctorate |
| Common to see ticks in Iquitos | Questionnaire | In the area where you live, is it common to see ticks in dogs? | No*<br>I do not know<br>Yes | No/I do not know*<br>Yes |
| Tick identification | Questionnaire | We showed the owner 4 vials with a tick larva, a nymph, a female adult, and a male adult, and ask | Did not identified any<br>Identified the larva<br>Identified the nymph<br>Identified at least one adult | Did not identified three stages*<br>Identified three stages |

| | | | | |
|---|---|---|---|---|
| | | him/her which one(s) were ticks. | | |
| Dogs can get TBDs[ϕ] | Questionnaire | Can dogs get diseases transmitted by ticks? | No*<br>I do not know<br>Yes | No/I do not know*<br>Yes |
| Know TBDs that affect dogs[ϕ] | Questionnaire | What TBD can dogs get? | Open question | No, owner cannot name any TBD that affect dogs*<br>Yes, owner can mention at least one TBD that affect dogs |
| Heard about Ehrlichia | Questionnaire | Have you ever heard about ehrlichiosis or Ehrlichia in dogs? | No<br>Yes | No*<br>Yes |
| Humans can get TBDs[ϕ] | Questionnaire | Can humans get diseases transmitted by ticks? | No*<br>I do not know<br>Yes | No/I do not know*<br>Yes |

*Reference categories in the regression analyses.
[γ]References: Age categories (Harvey, 2021), body condition score (American Animal Hospital Association, 2019)
**Observations from the visual inspection were registered in the home inspection form.
[ϕ] TBDs = Tick-borne diseases

**Sup. Table 2.** Primers sets and probes sequences for the detection of E. canis dsb gene and the IAC by qPCR

| Target | Probe/Primer | Sequence |
|---|---|---|
| *dbs* | Probe | FAM/BHQ1 5' -AGC TAG TGC TGC TTG GGC AAC TTT GAG TGA A-3' |
| | Forward | 5'-TTG CAA AAT GAT GTC TGA AGA TAT GAA ACA-3' |
| | Reverse | 5'-GCT GCA CCA CCG ATA AAT GTA TCC CCT A-3' |
| IAC | Probe | VIC/ MGB 5'-AGC ATC TGT TCT TGA AGG T-3’ |
| | Forward | 5'-ACC GTC ATG GAA CAG CAC GTA 3' |
| | Reverse | 5'-CTC CCG CAA CAA ACC CTA TAA AT-3' |

**Sup. Table 3.** Primers sequences for the detection of *E. canis* trp36 gene by PCR

| Target | Probe/Primer | Sequence |
|---|---|---|
| *trp36* | Forward | 5′-TTTAAAACAAAATTAACACACTA-3′ |
| | Reverse | 5′-AAGATTAACTTAATACTCAATATTACT-3 |

**Sup. Table 4.** Comparison of dog-, owner-, and home-related factors between individuals who did not indicate their home income and individuals who did indicate their home income

| | Total[⊥] | Missing home income | | p-value* |
|---|---|---|---|---|
| | | No (257, 90.2%) | Yes (28, 9.8%) | |
| **Dog-related factors**** | | | | |
| **Sex** | | | | 0.780 |
| Female | 119 (41.8%) | 108 (90.8%) | 11 (9.2%) | |
| Male | 166 (58.2%) | 149 (89.8%) | 17 (10.2%) | |
| **Neuter/Spayed** | | | | 0.937 |
| No | 242 (85.2%) | 218 (90.1%) | 24 (9.9%) | |
| Yes | 42 (14.8%) | 38 (90.5%) | 4 (9.5%) | |
| **Age** | | | | 0.187 |
| Junior | 98 (34.4%) | 92 (93.9%) | 6 (6.1%) | |
| Adult | 127 (44.6%) | 114 (89.8%) | 13 (10.2%) | |
| Senior | 60 (21.1%) | 51 (85%) | 9 (15%) | |
| **Body condition** | | | | 0.092 |
| Very thin | 4 (1.4%) | 3 (75%) | 1 (25%) | |
| Thin | 67 (23.5%) | 56 (83.6%) | 11 (16.4%) | |
| Ideal | 187 (65.6%) | 174 (93%) | 13 (7%) | |
| Overweight | 24 (8.4%) | 22 (91.7%) | 2 (8.3%) | |
| Obesity | 3 (1.1%) | 2 (66.7%) | 1 (33.3%) | |
| **Lifestyle** | | | | 0.393 |
| Only indoors | 113 (39.7%) | 104 (92%) | 9 (8%) | |
| Indoors/outdoors or only outdoors | 172 (60.3%) | 153 (89%) | 19 (11%) | |
| **Recent travel** | | | | 0.813 |
| No | 258 (90.5%) | 233 (90.3%) | 25 (9.7%) | |
| Yes | 27 (9.5%) | 24 (88.9%) | 3 (11.1%) | |
| **Current tick infestation** | | | | **0.024** |
| No | 139 (48.8%) | 131 (94.2%) | 8 (5.8%) | |
| Yes | 146 (51.2%) | 126 (86.3%) | 20 (13.7%) | |
| **History of tick infestation** | | | | 0.284 |
| No | 75 (26.3%) | 70 (93.3%) | 5 (6.7%) | |
| Yes | 210 (73.7%) | 187 (89%) | 23 (11%) | |
| **Frequently taken to vet** | | | | 0.241 |
| No | 162 (56.8%) | 149 (92%) | 13 (8%) | |
| Yes | 123 (43.2%) | 108 (87.8%) | 15 (12.2%) | |
| **Acaricide product used** | | | | 0.149 |
| No | 48 (16.8%) | 46 (95.8%) | 2 (4.2%) | |
| Yes | 237 (83.2%) | 211 (89%) | 26 (11%) | |
| **Home-related factors** | | | | |
| **District of residency** | | | | 0.304 |
| Iquitos | 96 (33.7%) | 88 (91.7%) | 8 (8.3%) | |
| San Juan | 95 (33.3%) | 86 (90.5%) | 9 (9.5%) | |
| Punchana | 47 (16.5%) | 39 (83%) | 8 (17%) | |
| Belén | 47 (16.5%) | 44 (93.6%) | 3 (6.4%) | |
| **Wall material**** | | | | |
| Brick or cement | | | | 0.987 |
| No | 40 (14.2%) | 36 (90%) | 4 (10%) | |
| Yes | 242 (85.8%) | 218 (90.1%) | 24 (9.9%) | |
| Plywood | | | | 0.812 |
| No | 135 (47.9%) | 121 (89.6%) | 14 (10.4%) | |

| | | | | |
|---|---|---|---|---|
| Yes | 147 (52.1%) | 133 (90.5%) | 14 (9.5%) | |
| Wood | | | | 0.446 |
| No | 152 (53.9%) | 135 (88.8%) | 17 (11.2%) | |
| Yes | 130 (46.1%) | 119 (91.5%) | 11 (8.5%) | |
| Corrugated iron | | | | **0.017** |
| No | 221 (78.4%) | 204 (92.3%) | 17 (7.7%) | |
| Yes | 61 (21.6%) | 50 (82%) | 11 (18%) | |
| **Indoor floor material**** | | | | |
| Cement | | | | 0.109 |
| No | 64 (22.8%) | 61 (95.3%) | 3 (4.7%) | |
| Yes | 217 (77.2%) | 192 (88.5%) | 25 (11.5%) | |
| Tile | | | | 0.498 |
| No | 195 (69.4%) | 174 (89.2%) | 21 (10.8%) | |
| Yes | 86 (30.6%) | 79 (91.9%) | 7 (8.1%) | |
| Dirt | | | | 0.584 |
| No | 222 (79%) | 201 (90.5%) | 21 (9.5%) | |
| Yes | 59 (21%) | 52 (88.1%) | 7 (11.9%) | |
| Wood | | | | 0.088 |
| No | 257 (91.5%) | 229 (89.1%) | 28 (10.9%) | |
| Yes | 24 (8.5%) | 24 (100%) | 0 | |
| **Number of dogs in home** | | | | 0.698 |
| One | 173 (60.7%) | 158 (91.3%) | 15 (8.7%) | |
| Two | 72 (25.3%) | 64 (88.9%) | 8 (11.1%) | |
| Three or more | 40 (14%) | 35 (87.5%) | 5 (12.5%) | |
| **Seeing ticks in home** | | | | **0.009** |
| No | 138 (48.4%) | 131 (94.9%) | 7 (5.1%) | |
| Yes | 147 (51.6%) | 126 (85.7%) | 21 (14.3%) | |
| **Owner-related factors** | | | | |
| **Level of education** | | | | 0.150 |
| No studies – less than high school | 34 (12%) | 31 (91.2%) | 3 (8.8%) | |
| High school (incomplete) | 24 (8.5%) | 22 (91.7%) | 2 (8.3%) | |
| High school (complete) | 78 (27.5%) | 67 (85.9%) | 11 (14.1%) | |
| Institute (incomplete) | 26 (9.2%) | 22 (84.6%) | 14 (15.4%) | |
| Institute (complete) | 46 (16.2%) | 43 (93.5%) | 3 (6.5%) | |
| University (incomplete) | 31 (10.9%) | 26 (83.9%) | 5 (16.1%) | |
| University (complete) - Graduate | 45 (15.9%) | 45 (100%) | 0 | |
| **Tick identification (all stages)** | | | | 0.894 |
| No | 147 (53.1%) | 133 (90.5%) | 14 (9.5%) | |
| Yes | 130 (46.9%) | 117 (90%) | 13 (10%) | |
| **Common to see ticks in dogs in Iquitos** | | | | 0.378 |
| No | 59 (20.7%) | 55 (93.2%) | 4 (6.8%) | |
| Yes | 226 (79.3%) | 202 (89.4%) | 24 (10.6%) | |
| **Dogs can get TBD** | | | | 0.666 |
| No | 43 (15.1%) | 38 (88.4%) | 5 (11.6%) | |
| Yes | 242 (84.9%) | 219 (90.5%) | 23 (9.5%) | |
| **Know TBD that affect dogs** | | | | 0.889 |
| No | 219 (90.5%) | 198 (90.4%) | 21 (9.6%) | |
| Yes | 23 (9.5%) | 21 (91.3%) | 2 (8.7%) | |
| **Heard about Ehrlichia** | | | | 0.880 |
| No | 237 (83.2%) | 214 (90.3%) | 23 (9.7%) | |
| Yes | 48 (16.8%) | 43 (89.6%) | 5 (10.4%) | |
| **Humans can get TBD** | | | | 0.356 |
| No | 90 (31.6%) | 79 (87.8%) | 11 (12.2%) | |

| | | | |
|---|---|---|---|
| Yes | 195 (68.4%) | 178 (91.3%) | 18 (8.7%) |

*p-value calculated using chi-square tests

**One dog was randomly selected per home to perform this analysis

### Sociodemographic survey questionnaire - English

#### Location

| | |
|---|---|
| Record ID | ______________________ |
| Interviewer: Home Code | ______________________ |
| Interviewer: Home Latitude | ______________________<br>(Collect by looking at the door of the home) |
| Interviewer: Home Longitude | ______________________<br>(Collect by looking at the door of the home) |
| Address | ______________________ |
| District | ______________________ |

### Overview

| | |
|---|---|
| How many people live in this home? | ______________________ |
| Does a dog live in this home? | ◯ No<br>◯ Yes |
| How many dogs live in this home? | ______________________ |
| Interviewer: Person over 18 years of age accepts participate in the study and signs informed consent | ◯ No<br>◯ Yes |
| Participant Code | ______________________ |
| Interview Date | ______________________ |
| Date of sample collection | ______________________ |
| Sample Collection Time | ______________________<br>(Indicate the time range from the beginning to the completion of the sample collection) |
| Phone number | ______________________ |
| Email | ______________________ |

### Sociodemographic information

#### First, I am going to ask you some questions about yourself and your family.

What is your full name?

__________________________________

What is your gender?

- ◯ Female
- ◯ Male
- ◯ I prefer not to say

How old are you?

__________________________________

What is the highest degree of education you have finished?

- ◯ No studies
- ◯ Initial
- ◯ Incomplete primary school
- ◯ Complete primary school
- ◯ Incomplete secondary school
- ◯ Complete secondary school
- ◯ Incomplete technical
- ◯ Complete technical
- ◯ Complete university
- ◯ Incomplete university
- ◯ Master's/doctoral degree

What kind of job do you have?

- ◯ State-dependent worker
- ◯ Dependent worker of a private company
- ◯ Self-employed worker

What is your primary occupation?

- ◯ Health Services
- ◯ Aquaculture/Fisheries
- ◯ Forestry/Logging/Timber
- ◯ Livestock
- ◯ Construction
- ◯ Transportation (motorcycle taxi, bus, small-small, or others)
- ◯ Domestic Worker
- ◯ Teacher
- ◯ Student
- ◯ Merchant
- ◯ Cook/Food Vendor
- ◯ Housewife
- ◯ Other

Specify your primary occupation (if Other)

__________________________________

The area where you work is:

- ◯ Rural
- ◯ Peri-urban
- ◯ Urban

What is your monthly (personal) income from this work?

- ◯ I do not generate income
- ◯ Less than or equal to S/. 500
- ◯ Between S/.501 and S/.1,000
- ◯ Between S/.1,001 and S/.1,500
- ◯ Between S/.1,501 and S/.2,000
- ◯ Between S/.2,001 and S/.2,500
- ◯ More than 2,500

Number of inhabitants contributing to home income ______________________

What is your total monthly home income?

- ◯ I don't generate income
- ◯ Less than or equal to S/.500
- ◯ Between S/.501 and S/.1,000
- ◯ Between S/.1,001 and S/.1,500
- ◯ Between S/.1,501 and S/.2,000
- ◯ Between S/.2,001 and S/.2,500
- ◯ Over 2,500

In this district, are you a resident or visitor?

- ◯ Visitor
- ◯ Resident

How long have you lived in this district?

______________________
(Indicate time in years and months)

### Characteristics of the home

#### Now I'm going to ask you some questions about your home.

Does your home receive electricity through a public grid?
- ◯ No
- ◯ Yes

In your home, what is the main source of water?
- ◯ Public system, exclusive use of family
- ◯ Public system, shared with other homes
- ◯ Public pipe, fountain outside the home, shared with the community
- ◯ Well water
- ◯ River, lake or canal water
- ◯ Truck/water tank
- ◯ Other

Specify the water source (if Other) __________________________________

The bathroom you have in your home is connected to:
- ◯ Public drainage network inside the home
- ◯ Public drainage network outside the home
- ◯ Septic latrine
- ◯ Latrine
- ◯ River, ditch or irrigation canal
- ◯ No bathroom (open field)
- ◯ Other

Specify what your bathroom is connected to (if Other) __________________________________

Does your home have a patio/orchard?
- ◯ No
- ◯ Yes

#### Are there sources of pollution near your home?

| | No | Yes |
|---|---|---|
| Dumps | ◯ | ◯ |
| Abandoned homes | ◯ | ◯ |
| Uncultivated lots | ◯ | ◯ |
| Contaminated water | ◯ | ◯ |
| Open Drain | ◯ | ◯ |
| Other | ◯ | ◯ |

Specify the source of contamination (if Other) __________________________________

In the last year, have you witnessed stray animals near your home?
- ◯ No
- ◯ Yes

What stray animals have you witnessed near your home in the last year?
- ☐ Dogs
- ☐ Cats
- ☐ Other

Specify which stray animals you have witnessed near your home in the last year (if Other) __________________________________

### Experiences, knowledge and vulnerability about ticks

**Now I am going to ask you a few questions about your experiences and knowledge about ticks**

| | |
|---|---|
| First, I am going to show you some tubes and you will tell me which one or which ones contain ticks | ☐ I do not identify any<br>☐ I identify the larva<br>☐ I identify the nymph<br>☐ I identify the adult |
| Have you ever observed ticks in your home? | ◯ No<br>◯ Yes |
| When was the last time you observed a tick in your home? | ______________________ |
| How often do you see ticks in your home? | ◯ Rarely<br>◯ Sometimes<br>◯ Often<br>◯ Always |
| Where in the home do you see ticks?<br><br>You can check more than one option. | ☐ Walls<br>☐ Floor<br>☐ Ceiling<br>☐ Patio<br>☐ Facade<br>☐ Other |
| Specify where you see ticks (if Other) | ______________________ |
| Can people contract tick-borne diseases? | ◯ No<br>◯ Yes<br>◯ I don't know |

**Now I'm going to ask you some questions about dogs.**

| | |
|---|---|
| In the area where you live, it is common to see ticks in dogs? | ◯ No<br>◯ Yes |
| Where do dogs get infested with ticks?<br><br>(This question can be rephrased as: where do ticks climb on dogs?) | ______________________ |
| Can dogs contract transmitted diseases? because of ticks? | ◯ No<br>◯ Yes |
| What tick-borne disease(s) can dogs get? | ______________________ |
| Have you ever heard of ehrlichiosis or ehrlichia in dogs? | ◯ No<br>◯ Yes |

**What symptoms do you think ehrlichiosis can cause in dogs?**

**I will read you all the options, and you will indicate if it the answer is yes, no, or I don’t know**

| | No | Yes | I don't know |
|---|---|---|---|
| High fever | ○ | ○ | ○ |
| Weight loss | ○ | ○ | ○ |
| Decay/Sadness | ○ | ○ | ○ |
| Bleeding / Bleeding | ○ | ○ | ○ |
| Vomiting or diarrhea | ○ | ○ | ○ |
| Other | ○ | ○ | ○ |

What other symptom(s) can dogs with ehrlichiosis have? ________________________________

### Animal - Epidemiological data (repeated for each animal)

Now I'm going to ask you some questions about THIS dog. Please respond with information only about this animal, not about any others you may have.

Dog Code

(Use the same code as the labels assigned to the sample tubes)

______________________________

What is the name of this dog?

______________________________

What breed is this dog?

______________________________

How old is this dog?

______________________________
(Indicate age of the dog in years)

How many months old is this dog?

______________________________
(Please indicate months if owner knows the exact age)

How many kg does this dog weigh approximately?

______________________________
(Indicate weight in kg)

Is this dog female or male?

- ◯ Female
- ◯ Male

Has this dog had puppies?

- ◯ No
- ◯ Yes
- ◯ I don't know

How many litters has this dog had?

______________________________

Has this dog been spayed?

- ◯ No
- ◯ Yes
- ◯ I don't know

Where was this dog born?

______________________________
(Indicate district, province, and department)

How long have you had this dog?

______________________________
(Indicate approximate time in years and months)

Where did you acquire this dog?

______________________________

Do you usually take this dog to the vet?

- ◯ No
- ◯ Yes

How often do you take this dog to the vet?

- ◯ Monthly
- ◯ Every 3 months
- ◯ Every 6 months
- ◯ Every year
- ◯ Other

Specify how often you take this dog to the vet (if Other)

__________________________________

When was the last time you took this dog to the vet?

__________________________________
(Please indicate approximate date)

In the last year, has this dog received any medication such as pills and/or injections?

- ◯ No
- ◯ Yes
- ◯ I don't know

What medication did the dog receive?

__________________________________________

**Do you use any of these products to eliminate ticks from this dog?**

**I will read you all the options, and you will indicate yes or no.**

| | No | Yes |
|---|---|---|
| Spray | ◯ | ◯ |
| Bathroom | ◯ | ◯ |
| Powder | ◯ | ◯ |
| Pipette | ◯ | ◯ |
| Tablets | ◯ | ◯ |
| Others | ◯ | ◯ |

What is the name of the spray you use on this dog?

__________________________________

How often do you use sprays on this dog?

- ◯ Monthly
- ◯ Every 3 months
- ◯ Every 6 months
- ◯ Every year
- ◯ Other

Specify how often you use sprays on this dog (if Other)

__________________________________

When was the last time you used this spray on this dog?

__________________________________

| | |
|---|---|
| How do you apply this spray to this dog?<br><br>(For example, dilute it in water, apply it directly, etc.) | __________________________________ |
| What is the name of the product you use to bathe this dog? | __________________________________ |
| How often do you bathe this dog with this product? | ◯ Monthly<br>◯ Every 3 months<br>◯ Every 6 months<br>◯ Every year<br>◯ Other |
| Please specify how often you bathe this dog with this product (if Other) | __________________________________ |
| When was the last time you bathed this dog with this product? | __________________________________ |
| How do you use the bath product on this dog?<br><br>(For example, dilute it in water, apply it directly, etc.) | __________________________________ |
| What is the name of the powder you use on this dog? | __________________________________ |
| How often do you use powder on this dog? | ◯ Monthly<br>◯ Every 3 months<br>◯ Every 6 months<br>◯ Every year<br>◯ Other |
| Specify how often you use powders on this dog (if Other) | __________________________________ |
| When was the last time you applied this powder on this dog? | __________________________________ |
| How do you use the powder on this dog?<br><br>(For example, dilute it in water, apply it directly, etc.) | __________________________________ |
| What is the name of the pill you use on this dog? | __________________________________ |
| How often you use pills in this dog? | ◯ Monthly<br>◯ Every 3 months<br>◯ Every 6 months<br>◯ Every year<br>◯ Other |

| | |
|---|---|
| Specify how often you use pills on this dog (if Other) | __________________________ |
| When was the last time you gave this pill to this dog? | __________________________ |
| How do you use the pill on this dog?<br><br>(For example, he dilutes it in water and then takes it, gives it directly, etc.) | __________________________ |
| What is the name of the pipette you use on this dog? | __________________________ |
| How often do you use pipettes in this dog? | ◯ Monthly<br>◯ Every 3 months<br>◯ Every 6 months<br>◯ Every year<br>◯ Other |
| Specify how often you use tick pipettes on this dog (if Other) | __________________________ |
| When was the last time you used a pipette on this dog? | __________________________ |
| How do you use the pipette on this dog?<br><br>(For example, he dilutes it in water and then takes it, gives it directly, etc.) | __________________________ |
| Specify the name of the tick remover/prevention product/substance you use on this dog | __________________________ |
| How often do you use this product/substance for ticks on this dog? | ◯ Monthly<br>◯ Every 3 months<br>◯ Every 6 months<br>◯ Every year<br>◯ Other |
| Please specify how often you use this tick product/substance on this dog (if Other) | __________________________ |
| When was the last time you applied this product/substance to this dog? | __________________________ |
| How do you use this other product/substance on this dog?<br><br>(For example, dilute it in water, apply it directly, etc.) | __________________________ |
| Why do you have this dog? | ☐ As a guardian<br>☐ As a pet<br>☐ Another reason |

Indicate why else you have this dog

__________________________________

Where do you keep this dog most of the time?

- ◯ Alone inside the home
- ◯ Inside the home and in the patio
- ◯ Only in the patio
- ◯ Only outside the home (does not include the patio)
- ◯ Inside and outside the home (does not include the patio)

Where does this dog sleep?

- ◯ Outside the home
- ◯ Inside the home

Please tell me where in the home the dog sleeps

__________________________________

Has this dog traveled recently?

(Include travel to any other district, province or department)

- ◯ No
- ◯ Yes

How long ago did this dog travel?

__________________________________
(Indicate whether it is days, weeks, months, or years)

Where did this dog travel?

__________________________________
(Indicate district, province and department)

Was this an urban or rural area?

__________________________________

In the last year, have you ever observed ticks in this dog?

- ◯ No
- ◯ Yes

Does this dog get ticks in any special place?

Interviewer: Examples can be given such as "in the forest, in terrales, etc."

__________________________________
(Please indicate location)

**Has this dog ever had any of the following diseases?**

**I am going to read you all the options and you are going to tell me yes, no, or you don't know.**

| | No | Yes | I don't know |
|---|---|---|---|
| Rickettiosis | ◯ | ◯ | ◯ |
| Ehrlichiosis | ◯ | ◯ | ◯ |
| Malaria | ◯ | ◯ | ◯ |
| Babesiosis | ◯ | ◯ | ◯ |
| Anaplasmosis | ◯ | ◯ | ◯ |

### Animal - Clinical data (repeated for each animal)

**In the last month, this dog has presented:**

**I will read you all the options and you will tell me yes or no.**

| | No | Yes |
|---|---|---|
| **Vomiting** | ◯ | ◯ |
| **Diarrhea** | ◯ | ◯ |
| **Bleeding** | ◯ | ◯ |
| **Weakness/Weakness** | ◯ | ◯ |
| **Weight Loss** | ◯ | ◯ |
| **Loss of Appetite** | ◯ | ◯ |
| **Paleness** | ◯ | ◯ |
| **Other** | ◯ | ◯ |

---

Specify where was the dog bleeding from

- ☐ Mouth
- ☐ Nose
- ☐ Stool
- ☐ Urine
- ☐ Other

---

Specify what other symptoms this dog had in the last month

________________________________

---

Indicate the color of the dog's mucous membranes

- ◯ Pale
- ◯ Pink/white
- ◯ Intense red
- ◯ Blue or purple

---

Indicate the dog's body condition

- ◯ Very thin
- ◯ Slim
- ◯ Ideal
- ◯ Overweight
- ◯ Obesity

---

Indicate the skinfold return time

________________________________
(Indicate time in seconds)

---

Indicate the number of ticks observed on this dog

- ◯ 0
- ◯ 1-10
- ◯ 11-20
- ◯ 21-30
- ◯ >30

#### Indicate the samples collected from this animal.

| | No | Yes |
|---|---|---|
| **Ticks** | ○ | ○ |
| **Purple top tube** | ○ | ○ |
| **Red top tube** | ○ | ○ |

### Home characteristics

#### Wall material

| | No | Yes |
|---|---|---|
| Brick or cement | ○ | ○ |
| Corrugated iron | ○ | ○ |
| Stone with mud | ○ | ○ |
| Wood | ○ | ○ |
| **Plywood/Mat** | ○ | ○ |
| Pona | ○ | ○ |
| Other | ○ | ○ |

Percentage of brick or cement ______________________

Percentage of corrugated iron ______________________

Percentage of stone or mud ______________________

Percentage of wood ______________________

Percentage of plywood/mat ______________________

Percentage of pona ______________________

Specify the type of wall material if "Other" was answered ______________________

Percentage of other material in walls ______________________

#### Floor material

| | No | Yes |
|---|---|---|
| Parquet or polished wood | ○ | ○ |
| Asphalt, vinyl sheets | ○ | ○ |
| Tiles, ceramic tiles | ○ | ○ |
| **Soil** | ○ | ○ |
| Wood | ○ | ○ |
| Cement | ○ | ○ |
| Pona | ○ | ○ |
| Other | ○ | ○ |

| | |
|---|---|
| Percentage of parquet or polished wood | __________________________________ |
| Percentage of asphalt sheets, vinyls | __________________________________ |
| Percentage of tiles | __________________________________ |
| Percentage of soil | __________________________________ |
| Percentage of wood | __________________________________ |
| Percentage of cement | __________________________________ |
| Specify the type of flooring material if "Other" was answered | __________________________________ |
| Percentage of other material | __________________________________ |
| Does the home have a patio? | ◯ No<br>◯ Yes |
| Does the home have an orchard? | ◯ No<br>◯ Yes |
| Patio (outside the home): Presence of accumulated garbage/unusable materials | ◯ No<br>◯ Yes |
| Orchard (inside the home): Presence of accumulated garbage/unusable materials | ◯ No<br>◯ Yes |

#### Patio (outside the home): Presence of a water body.

| | No | Yes |
|---|---|---|
| Open Drain | ◯ | ◯ |
| Standing water | ◯ | ◯ |
| Cocha | ◯ | ◯ |
| Artesian well | ◯ | ◯ |

#### Orchard (inside the home): Presence of a water body.

| | No | Yes |
|---|---|---|
| Open drain Pond | ◯ | ◯ |
| water Cocha | ◯ | ◯ |
| Artesian well | ◯ | ◯ |

#### Patio (outside the home): Type of soil

| | 0 | 1-10 | 11-20 | 21-30 | 31-40 | 41-50 | 51-60 | 61-70 | 71-80 | 81-90 | 91-100 |
|---|---|---|---|---|---|---|---|---|---|---|---|
| Brick or cement | ◯ | ◯ | ◯ | ◯ | ◯ | ◯ | ◯ | ◯ | ◯ | ◯ | ◯ |
| Earth | ◯ | ◯ | ◯ | ◯ | ◯ | ◯ | ◯ | ◯ | ◯ | ◯ | ◯ |
| **Grass/plants/grass** | ◯ | ◯ | ◯ | ◯ | ◯ | ◯ | ◯ | ◯ | ◯ | ◯ | ◯ |
| Sand | ◯ | ◯ | ◯ | ◯ | ◯ | ◯ | ◯ | ◯ | ◯ | ◯ | ◯ |
| Pona | ◯ | ◯ | ◯ | ◯ | ◯ | ◯ | ◯ | ◯ | ◯ | ◯ | ◯ |
| Madera | ◯ | ◯ | ◯ | ◯ | ◯ | ◯ | ◯ | ◯ | ◯ | ◯ | ◯ |

#### Orchard (inside the home): Type of soil

| | 0 | 1-10 | 11-20 | 21-30 | 31-40 | 41-50 | 51-60 | 61-70 | 71-80 | 81-90 | 91-100 |
|---|---|---|---|---|---|---|---|---|---|---|---|
| Brick or Cement Soil | ◯ | ◯ | ◯ | ◯ | ◯ | ◯ | ◯ | ◯ | ◯ | ◯ | ◯ |
| **Grass/Plants/Grass** | ◯ | ◯ | ◯ | ◯ | ◯ | ◯ | ◯ | ◯ | ◯ | ◯ | ◯ |
| Sand | ◯ | ◯ | ◯ | ◯ | ◯ | ◯ | ◯ | ◯ | ◯ | ◯ | ◯ |
| Pona | ◯ | ◯ | ◯ | ◯ | ◯ | ◯ | ◯ | ◯ | ◯ | ◯ | ◯ |
| Madera | ◯ | ◯ | ◯ | ◯ | ◯ | ◯ | ◯ | ◯ | ◯ | ◯ | ◯ |
| | ◯ | ◯ | ◯ | ◯ | ◯ | ◯ | ◯ | ◯ | ◯ | ◯ | ◯ |

Type of access to home

◯ Paved Street
◯ Dirt Street
◯ Other

#### Types of boundaries of the property/home.

| | No | Yes |
|---|---|---|
| Vegetation boundary | ◯ | ◯ |
| Wooden border | ◯ | ◯ |
| Wire boundary | ◯ | ◯ |
| Concrete boundary | ◯ | ◯ |
| No boundary | ◯ | ◯ |

Percentage of vegetation boundary ______________________

Percentage of the wooden boundary ______________________

Percentage of wire boundary ______________________

Percentage of concrete boundary ______________________

| | |
|---|---|
| Specify the type of access to the home (if Other) | __________________________________ |
| Daytime shade level within a radius of < 10 m | __________________________________ (Take into account that the shade permanently covers the aforementioned radius during the day) |
